# Host transcriptional response to TB preventive therapy differentiates two sub-groups of IGRA-positive individuals

**DOI:** 10.1101/2020.07.20.202986

**Authors:** Claire Broderick, Jacqueline M Cliff, Ji-Sook Lee, Myrsini Kaforou, David AJ Moore

## Abstract

We investigated the longitudinal whole blood transcriptional profile responses to tuberculosis preventive therapy of 18 IGRA-positive tuberculosis contacts and IGRA-negative, tuberculosis-unexposed healthy controls.

Longitudinal unsupervised clustering analysis with a subset of 474 most variable genes in antigen-stimulated blood separated the IGRA+ participants into two distinct subgroups, one of which clustered with the IGRA-negative controls. 117 probes were significantly differentially expressed over time between the two cluster groups, many of them associated with immunological pathways important in mycobacterial control.

We contend that the differential host RNA response reflects lack of *M.tuberculosis* (*Mtb*) viability in the group that clustered with the IGRA-unexposed healthy controls, and *Mtb* viability in the group (1/3 of IGRA-positives) that clustered away.

Gene expression patterns in the blood of IGRA+ individuals emerging during the course of PT, which reflect *Mtb* viability, could have major implications in the identification of risk of progression, treatment stratification and biomarker development.

## Introduction

The term latent tuberculosis infection (LTBI) is loaded with the inference that viable *Mycobacterium tuberculosis* (*Mtb*) organisms are present in the affected individual which, under the right circumstances, have the capacity to cause reactivation and TB disease. Tests of immunological reactivity, whether delayed type hypersensitivity reactions measured in the tuberculin skin test (TST) or T lymphocyte stimulation though antigen recognition in the interferon gamma release assays (IGRAs) are widely referred to as tests for LTBI [1].

However, neither approach demonstrates presence of viable *Mtb* bacilli and there is no histopathological hallmark of LTBI. The lifetime risk of reactivation disease from an *Mtb* infection acquired remotely in time is around 10%, with most of that risk believed to arise during the first five years after infection [2]. In the interval between acquisition of infection and development of disease, *Mtb* maintains viability and is assumed to be slowly replicating, either under close immunological control or in a relatively immunologically privileged location. Thus LTBI induces immunological sensitization as reflected in the TST and IGRA, tests that demonstrate immunological memory for prior exposure to mycobacterial antigens.

Nevertheless, 90% of individuals demonstrating immunological recognition of *Mtb* antigens by positive IGRA or TST never develop active TB disease. Taking the inherent assumption that TST and IGRA are indicators of LTBI to its logical conclusion, the 90% who escape development of TB do so because the immune control-pathogen balance remains in favor of the human host. An alternative explanation might be that a large proportion of those with positive TST and IGRA testing do not harbor viable organisms and are thus incapable of progressing to reactivation TB.

Preventive therapy (PT), in which a limited course of anti-TB antibiotics is used to sterilize presumed viable infection in individuals with positive TST and/or IGRA tests, has been shown to be highly effective in reducing the risk of future TB disease [3].

We hypothesized that differentiation of LTBI with viable bacilli from immunological sensitization without viable infection could be achieved by investigating the whole blood transcriptomic response to effective PT. We hypothesized that mycobacterial killing from effective LTBI PT would lead to a detectable alteration in the transcriptome that would not be seen in those individuals in whom there were no *Mtb* to be killed, whether these were IGRA/TST positive or healthy IGRA/TST negative controls with no known prior TB exposure.

## Results

### Recruitment of participants

Thirty adult IGRA-positive (IGRA+) participants were recruited to the study in the period October 2016 to January 2018, of whom 20 took a 12-week course of daily combined rifampicin/ isoniazid (RH) as preventive therapy (PT) and completed study follow-up. Adult IGRA-negative (IGRA-) healthy volunteers were recruited to the study and completed a two-week course of daily RH. Blood samples were collected from all participants at baseline (V1) and 2 weeks after initiating RH (V2), with an additional sample point in IGRA+ participants within 6 weeks of completion of the 12-week course of treatment (V3). At every timepoint, an unstimulated PAXgene whole blood sample and a stimulated blood sample (via QuantiFERON TB Gold Plus, Qiagen) was collected.

After quality control and pre-processing, 18 IGRA+ individuals and 4 IGRA-healthy controls were taken forward for comparator analyses (Figure 1, Figure 1-figure supplement 1). Recent exposure to drug-susceptible pulmonary TB was confirmed for 15/18 IGRA+s. There was no significant difference in age, gender, ethnicity or BCG status between the 18 IGRA+s and 4 IGRA-healthy controls (Table 1).

**Figure 1.**
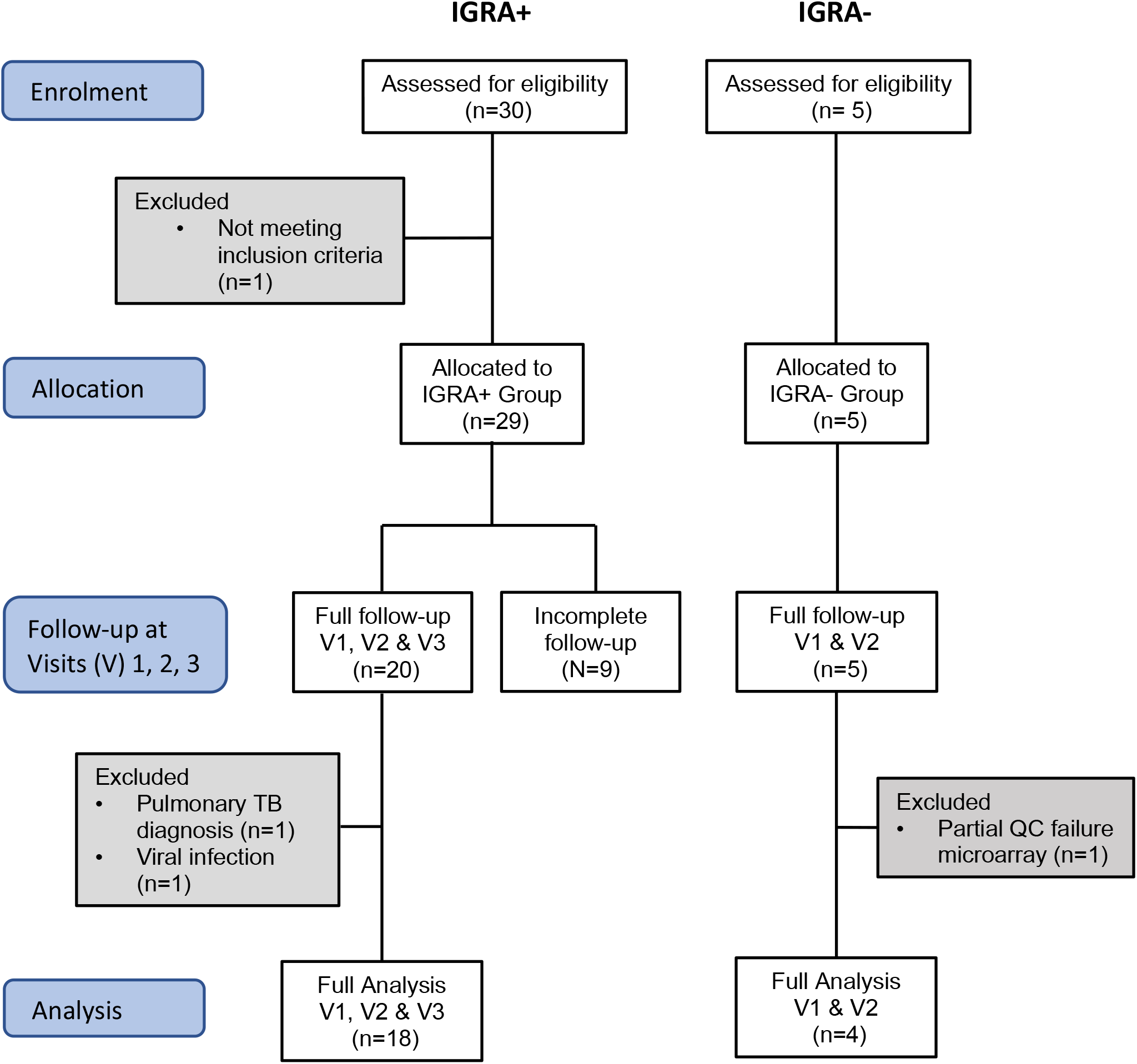
Study overview, showing patient numbers and exclusions.

**Table 1.** Subject Characteristics.

|  |  | IGRA+ group | IGRA- Healthy control group |
| --- | --- | --- | --- |
| Number |  | 18 | 4 |
| Age in years: Median (IQR) |  | 34 (28-38) | 28 (27-29) |
| Gender | Male | 10 (56%) | 3 (75%) |
|  | Female | 8 (44%) | 1 (25%) |
| Confirmed recent drug-susceptible TB exposure | Yes | 15 (83%) | 0 (0%) |
|  | No | 3 (17%) | 4 (100%) |
| BCG | Yes | 14 (78%) | 2 (50%) |
|  | No | 2 (11%) | 2 (50%) |
|  | Unknown | 2 (11%) | 0 (0%) |
| Continent of Birth | Africa | 4 (22%) | 0 (0%) |
|  | Asia | 4 (22%) | 0 (0%) |
|  | Australasia | 0 (0%) | 1 (25%) |
|  | Europe | 9 (50%) | 2 (50%) |
|  | North America | 0 (0%) | 1 (25%) |
|  | South America | 1 (6%) | 0 (0%) |
|  | Unknown | 0 (0%) | 0 (0%) |
| Ethnicity | Asian<sup>1</sup> | 5 (28%) | 2 (50%) |
|  | Black<sup>2</sup> | 4 (22%) | 0 (0%) |
|  | White<sup>3</sup> | 8 (44%) | 2 (50%) |
|  | Other<sup>4</sup> | 1 (6%) | 0 (0%) |

<sup>1</sup>Includes Bengali, Hong Kong, Kurdish, Sri Lankan, Turkish; <sup>2</sup>Includes Black African; <sup>3</sup>Includes White British, Polish, Romanian, White other; <sup>4</sup>Includes Latin American, Unknown

### Comparing gene expression profiles for IGRA+ versus IGRA-participants

First, we evaluated whether there were discernable differences in gene expression between the IGRA+ participants and IGRA-healthy controls, using linear models[4]. In the unstimulated PAXgene blood samples, no transcripts were found to be significantly differentially expressed (SDE) between the IGRA+ and IGRA-participants at baseline (V1) or V2 (Benjamini-Hochberg [BH] corrected p value < 0.05).

In this study, QuanitFERON-TB Gold Plus TB1 and TB2 tubes were used to stimulate whole blood. While both tubes contain peptides from ESAT-6 and CFP-10 *Mycobacterium tuberculosis* (*Mtb*) antigens, the TB1 tube peptides are designed to stimulate CD4+ T cells, and the TB2 peptides to stimulate both CD4+ and CD8+ T cells [5]. In contrast to the PAXgene tube whole blood samples, in the TB1-stimulated samples, 123 transcripts were SDE between IGRA+ and IGRA-individuals in the baseline (V1) samples and 93 were SDE between IGRA+ and IGRA-individuals in the V2 samples (BH corrected p value < 0.05) (Figure 2A and 2B and listed in Supplementary File 1). In the TB2-stimulated blood samples, when IGRA+ individuals were compared to IGRA-, 43 transcripts were found to be SDE in the V1 samples and 86 in the V2 samples. (BH corrected p value < 0.05) (Figure 2C and 2D and listed in Supplementary File 1). In summary, in vitro stimulation was necessary to distinguish the IGRA+ group from the IGRA-group.

**Figure 2.**
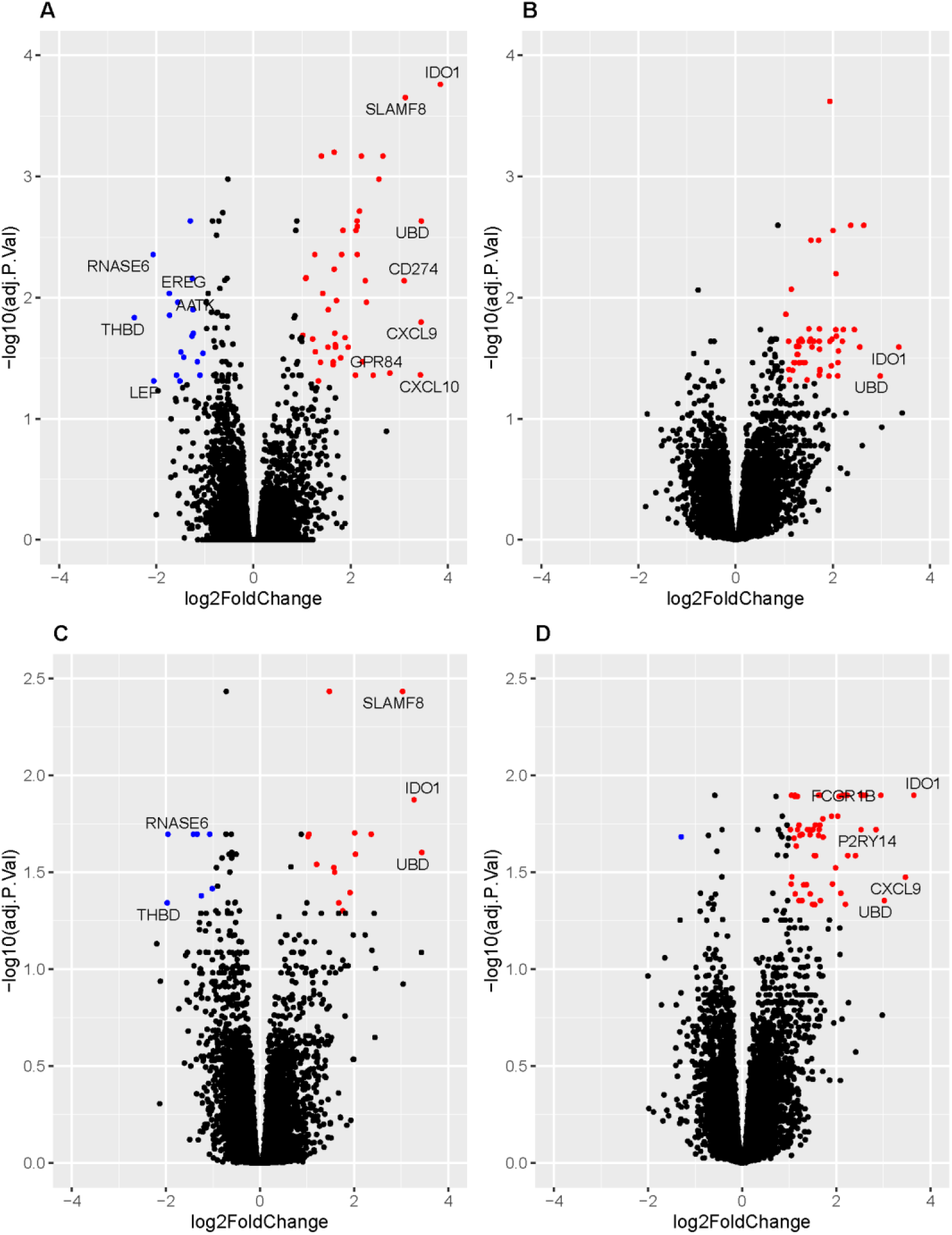
Volcano plots showing genes significantly differentially expressed between IGRA+ and IGRA-individuals. Genes upregulated in IGRA+s with log2Foldchange (LFC) >1 and Benjamini-Hochberg adjusted p value <0.05 are shown in red. Genes downregulated in IGRA+ individuals with LFC <-1 and BH adjusted p value <0.05 are shown in blue. Genes with LFC >2.7 and < −1.7 are annotated with their gene symbols. Plots are shown for TB1-stimulated samples at Visit (V) 1 [A] and V2 [B] and TB2-stimulated samples at V1 [C] and V2 [D]

### Effects of stimulation on whole blood gene expression

In addition to the TB1 and TB2 *Mtb*-peptide-containing tubes, the QuantiFERON-TB Gold Plus kit also includes a “negative” tube which contains no mycobacterial antigen peptides We assessed the effects of stimulation by comparing gene expression in the TB1- and TB2-stimulated tubes versus the negative tube at visit 1, using paired t-tests. In the IGRA+ group, when TB1 tube samples were compared to the negative tube, 3578 transcripts were SDE, while 3217 transcripts were SDE in the TB2 tube samples vs. the negative tube samples (BH corrected p value < 0.05), 2495 of which overlapped with the TB1 comparison (Figure 2-figure supplement 1A and 1B; SDE transcripts listed in Supplementary File 2). No genes were found to be SDE for the TB1-vs TB2-stimulated samples comparison.

In the IGRA-healthy controls, 37 transcripts were SDE in the TB1-stimulated samples compared to the negative tubes at visit 1 whereas just four transcripts were SDE in the TB2-stimulated samples (BH corrected p value < 0.05) (Figure 2- figure supplement 1C and 1D; SDE transcripts listed in Supplementary File 3).

### Filtering the gene expression dataset

Analyses were focused on the stimulated samples, as there had been no detectable differences between the IGRA+ and IGRA-participants in the unstimulated PAXgene samples. As described above, stimulation induced changes in gene expression in the IGRA-healthy controls, with a higher number of SDE genes observed with TB1-stimulation than TB2-stimulation, suggesting a greater non-specific effect independent of *Mtb* infection in the TB1 stimulation. Therefore, we focused on the TB2-stimulated samples for the next stage of the analysis.

The gene set was filtered to eliminate noise. Those genes that were lowly expressed or with extreme outlying values were removed, and of the remaining transcripts, those with the greatest variability between participants and over time were selected for the analysis, with X-transcripts SDE with gender and Y-chromosome transcripts removed. Through this process, a dataset with the “most variable genes” was generated for the TB2-stimulated samples (474 transcripts, listed in Supplementary File 4).

### Clustering analysis of longitudinal gene expression

We hypothesized that the IGRA+ group is heterogeneous, containing individuals that would demonstrate a transcriptomic response to PT (those with viable mycobacteria), and IGRA+ individuals without viable mycobacteria, who would not demonstrate a transcriptomic response to PT and would more closely resemble the healthy control IGRA-group. To unmask the PT-specific transcriptomic responses, we sought to stratify the IGRA+ group of individuals in an agnostic way. We employed unsupervised clustering analysis of longitudinal gene expression in the 18 IGRA+ patients and the 4 IGRA-controls, aiming to identify IGRA+ subgroups, using the most variable 474 transcripts in the TB2-stimulated dataset. The BClustLong package in ‘R’ [6] was utilized, which uses a linear mixed-effects framework to model the trajectory of genes over time and bases clustering on the regression coefficients obtained from all genes.

This longitudinal clustering analysis revealed two subgroups of IGRA+ participants. One subgroup of IGRA+s (IGRA+ subgroup A, N=12) clustered with the four healthy controls (Cluster 1), suggesting their gene expression over time was more similar to this *Mtb*-unexposed IGRA-population than it was to the remaining IGRA+s (IGRA+ subgroup B, N=6) who formed cluster 2. There were no significant differences in age, gender, ethnicity, BCG vaccination status or the IGRA+ participants’ TB contact history between clusters 1 and 2 (Table 2).

**Table 2.** Characteristics of Cluster groups 1 and 2.

|  |  | BClustLong clustering group |  |  |
| --- | --- | --- | --- | --- |
|  |  | Cluster 1 | Cluster 2 | p value |
| Number of participants |  | 16 | 6 | N/A |
| Patient IDs |  | HC51<br>HC53<br>HC54<br>HC55<br>LTBI1<br>LTBI2<br>LTBI3<br>LTBI5<br>LTBI7<br>LTBI9<br>LTBI12<br>LTBI15<br>LTBI16<br>LTBI27<br>LTBI28 | LTBI6<br>LTBI10<br>LTBI14<br>LTBI22<br>LTBI23<br>LTBI30 | N/A |
| Age in years: Median (IQR) |  | 32.5 (24-41) | 33.5 (29-38) | 0.6 |
| Gender | Male | 9 (56%) | 4 (66%) | 1 |
|  | Female | 7 (44%) | 2 (33%) |  |
| Confirmed recent exposure to DS-TB<sup>1</sup> | Yes | 10 (83%) | 5 (83%) | 1 |
|  | No | 2 (17%) | 1 (17%) |  |
| BCG | Yes | 10 (62%) | 6 (100%) | 0.2 |
|  | No | 4(25%) | 0 (0%) |  |
|  | Unknown | 2 (13%) | 0 (0%) |  |
| Continent of Birth | Africa | 3 (19%) | 1 (17%) | 0.2 |
|  | Asia | 1 (6%) | 3 (50%) |  |
|  | Australasia | 1 (6%) | 0 (0%) |  |
|  | Europe | 9 (56%) | 2 (33%) |  |
|  | North America | 1(6%) | 0 (0%) |  |
|  | South America | 1 (6%) | 0 (0%) |  |
| Ethnicity | Asian<sup>2</sup> | 4 (25%) | 3 (50%) | 0.7 |
|  | Black<sup>3</sup> | 3 (19%) | 1 (17%) |  |
|  | White<sup>4</sup> | 8 (50%) | 2 (33%) |  |
|  | Other<sup>5</sup> | 1 (6%) | 0 (0%) |  |

<sup>1</sup>for IGRA+ participants only
<sup>2</sup>Includes Bengali, Hong Kong, Kurdish, Sri Lankan, Turkish; <sup>3</sup>Includes Black African; <sup>4</sup>Includes White British, Polish, Romanian, White other; <sup>5</sup>Includes Latin American, Unknown

### Longitudinal differential gene expression analysis

In order to unravel the underlying blood transcriptomic differences between the two cluster groups generated by the unsupervised clustering, we performed longitudinal differential gene expression analysis using MaSigPro package in R [7]. MaSigPro uses a two-step regression strategy to firstly identify genes with significant temporal expression changes and then identify those genes which are significantly differentially expressed between groups.

Of the 474 transcripts in the dataset, 117 transcripts corresponding to 109 genes, were SDE over time between the two patient groups (with degrees of freedom=1 capturing linear trends, BH corrected p value < 0.05, listed in Supplementary File 5), while 2 of these genes had significant linear terms associated with time (*P2RY6, SLC2A3*). Setting the degrees of freedom to 2, 69 out of the 117 genes were SDE over time between the two cluster groups (BH corrected p value < 0.05, listed in Supplementary File 5), while 4 of these genes (*MSR1, MT1CP, IGHG3, IGHG1*) had significant linear and quadratic terms associated with time as well. In comparing cluster 1 vs. cluster 2, when one of the clusters is heterogeneous (IGRA+ subgroup A plus IGRA-healthy controls), it is expected that some of the differences will be down to the IGRA+ subgroup B versus IGRA- and not the IGRA+ subgroup B vs IGRA+ subgroup A comparison.

These 109 genes largely encoded proteins with known immune system function. Around one quarter have been previously reported in transcriptomics studies comparing blood from TB patients with healthy controls (31 transcripts, 25 genes) or with other diseases (9 transcripts, 7 genes) [8–14]; (Supplementary File 5).

Coefficients obtained using MaSigPro were used to cluster significant genes with similar longitudinal expression patterns (Figure 3). Often the proteins contained within a gene set had similar function, such as the CXC chemokines CXCL9, 10 and 11 in gene set 2 which were more highly expressed in patient cluster 2 and increased at V2, and the pro-inflammatory NF-κB transcription factor-inducing proteins IFNγ, IL-1R associated kinase 2 (IRAK2) and TNF superfamily member 15 (TNFSF15) in gene set 4, which were more highly expressed in patient cluster 2 and decreased through PT. BATF2, GCH1 and GBP3 all grouped in gene set 9, with consistently higher expression in patient cluster 2. Gene expression was higher in patient cluster 1 in only one gene set (gene set 3).

**Figure 3.**
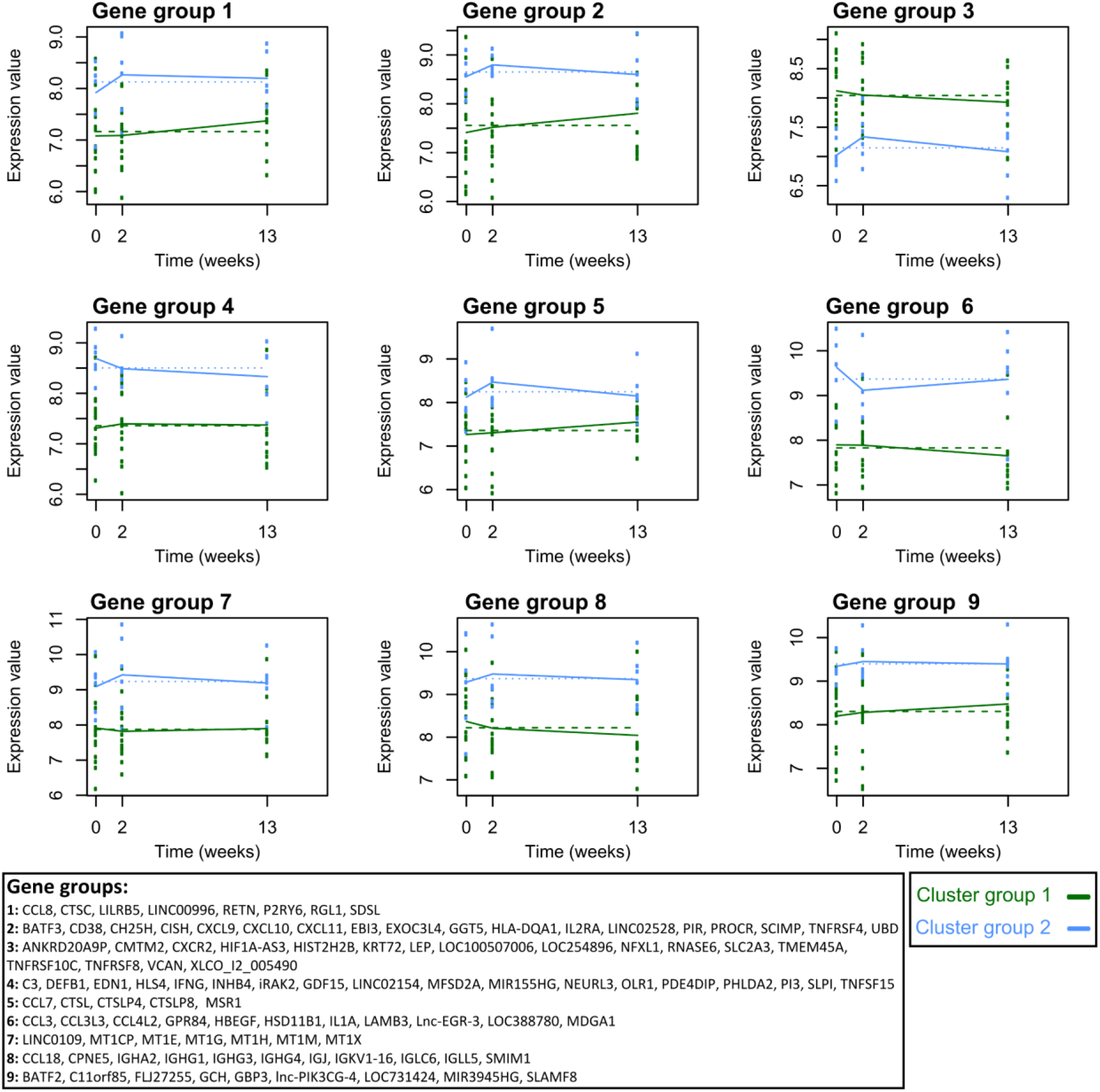
Longitudinal differential gene expression analysis between patient cluster groups 1 and 2 was performed using the TB2-stimulated whole blood samples. With 1 degree of freedom, 117/474 transcripts were SDE over time and between cluster groups 1 and 2 (BH corrected p value < 0.05). The coefficients obtained were used to group together significant genes with similar longitudinal expression patterns. MaSigPro identified 9 gene groups. Plots of gene expression against time for these gene groups are shown for patient cluster groups 1 (green) and 2 (blue). Lines join the median expression values of the gene groups at each timepoint. The gene symbols are listed for each gene group.

### Biological relevance of the significantly differentially expressed genes

The biological relevance of the 117 transcripts significantly differentially expressed over time between the two patient cluster groups was investigated. Biological pathways analysis was performed using Reactome pathway knowledgebase [15], with 80/117 transcripts successfully mapping to the database. Eleven pathways had significant over-representation of transcripts within our dataset (BH corrected p value < 0.05; listed in Supplementary file 6): these were all related to the immune system and encompassed pathways related to chemokine receptor binding, cytokine signaling – including IL10, TNF and regulatory T cells, metal ion binding and Complement cascade activation. There were a further 39 pathways with borderline over-representation: these largely encompassed biological functions related to innate immunity, antimicrobial peptides, phagocytosis, intracellular infection, and further cytokine signaling and Complement activation pathways.

### Differing cellular responses to preventive therapy

Relative cellular abundances were estimated from the gene expression data using CibersortX [16]. The estimated abundances of monocytes and lymphocytes were used to calculate the monocyte: lymphocyte ratio (MLR) for the two cluster groups at all three visits. At visits 1 and 3, the MLRs were similar between Clusters 1 and 2. However, at Visit 2, they were higher in cluster 2 (median= 0.52) compared to cluster 1 (median= 0.29, p=0.03). This difference at Visit 2 remained when the IGRA-healthy controls were removed from the analysis, with the MLR higher in IGRA+ subgroup B (median=0.52) compared to subgroup A (median=0.35, p=0.04) (Figure 4A).

**Figure 4.**
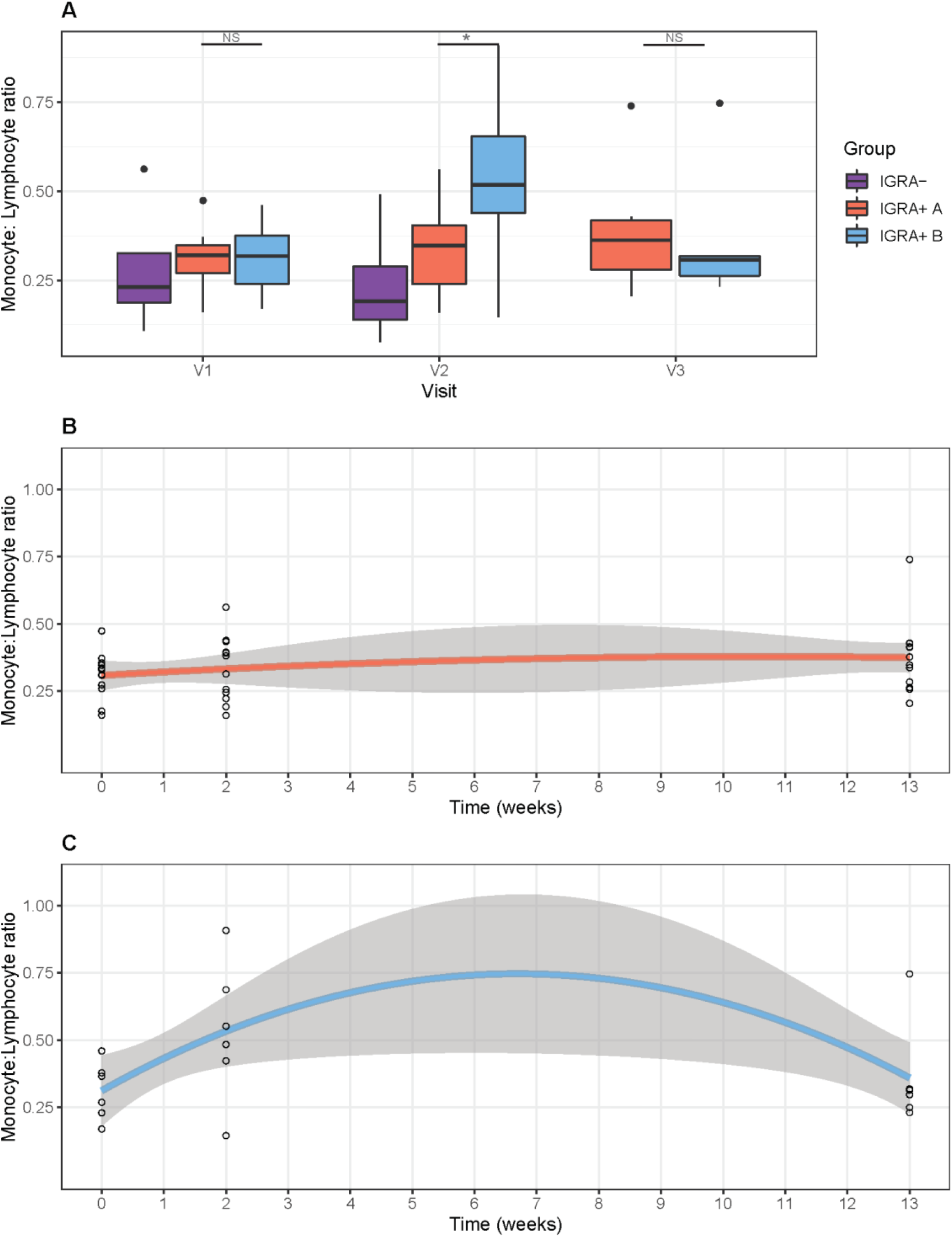
Cibersortx was used to estimate the abundance of monocytes and lymphocytes in the TB2-stimulated whole blood samples at each visit, and the monocyte: lymphocyte ratio was calculated. (A) Boxplots showing the Monocyte: Lymphocyte ratios at Visits 1, 2 and 3 for IGRA-healthy controls and IGRA+ groups A and B. NS denotes p >0.05, * denotes p≤0.05. Scatterplots showing the change in Monocyte: lymphocyte ratio over the time-course of the study period for (B) IGRA+ subgroup A and (C) IGRA+ subgroup B, where Visit 1 is 0 weeks, Visit 2 is 2 weeks and Visit 3 is 13 weeks, with 90% confidence intervals shown.

Using a second-degree polynomial model, the MLR was found to change over the time-course of the study period in IGRA+ subgroup B, and was close to the threshold of significance (linear term p=0.07, quadratic term p=0.06). This was not observed in IGRA+ subgroup A (linear term p=0.6, quadratic term p=0.8) (Figure 4B and C).

The relative abundances of other cell types including total monocytes, total lymphocytes, total CD4+ T cells and neutrophils were also observed to change with time in IGRA+ subgroup B and not subgroup A (Figure 4 –figure supplement 1).

## Discussion

This analysis has demonstrated that IGRA+ participants could be stratified according to their whole blood transcriptome into two distinct populations, one of which clustered with IGRA-, tuberculosis (TB)-unexposed controls. This separation was not clearly discernible when the transcriptomes of participants were evaluated at baseline in unstimulated whole blood, but rather was unmasked by TB-specific peptide stimulation after 14 days of TB preventive therapy (PT).

We hypothesized that PT would mediate mycobacterial death in participants for whom IGRA positivity was attributable to ongoing viable *Mycobacterium tuberculosis* (*Mtb*) infection and that the resulting immunological response, detected as a whole blood transcriptomic readout, would differentiate such individuals from a group of IGRA+ participants in whom PT would have no anti-mycobacterial effect due to the absence of viable *Mtb*. Our agnostic clustering approach clustered all four IGRA-healthy controls with a subgroup of IGRA+s (IGRA+ A), which is strongly suggestive that if indeed these clusters do define *Mtb* viability status then the true latent tuberculosis infection (LTBI) participants lie within the other subgroup (IGRA+ B). The genes differentially expressed between the two clusters through PT were predominantly involved in the immune system, particularly related to intracellular infection, inflammation, chemotaxis and cytokine signalling, indicating a biologically plausible specific response in the IGRA+ B subgroup.

Alternative explanations for the clear separation of these two groups were considered. Rifampicin has important antimicrobial effects against gram-positive organisms and can eliminate upper respiratory tract carriage of gram-negative organisms such as *Neisseria meningitidis* and *Haemophilus influenzae* within 2-4 days. The inclusion of rifampicin/isoniazid treated, IGRA-negative control participants was an attempt to capture and isolate any such non-mycobactericidal effect. In the absence of microbiological sampling and/or microbiome analysis we cannot entirely exclude the possibility that the separation of the groups is attributable to an effect completely unrelated to *Mtb* infection; however two factors which weigh against this alternative explanation are the low prevalence of *N. meningitidis* and *H. influenzae* carriage in this population (<10% combined) and the identification amongst the differentially expressed genes of several genes known to be associated with *Mtb* response pathways. The changes through PT overlapped with reported changes in blood transcriptome during treatment of active TB cases [17, 18]. The monocyte-to-lymphocyte ratio transiently increased only in the IGRA+ B subgroup: this ratio has been linked with TB disease susceptibility and blood transcriptomes [19]. The prevalence of carriage of non-tuberculous mycobacteria in this London-resident population would also be expected to be very low. Finally, we were concerned to exclude all possible artefactual explanations related to sample handling and found no effect association with study site, time to sample processing, study personnel or date of enrolment.

We contend that interferon gamma release assays (IGRA) and tuberculin skin tests (TST) are mis-represented as tests for LTBI, a term which infers viability of *Mtb* with potential to cause future reactivation disease. We believe that the observation that 90% of individuals with positive testing by IGRA/TST do not develop TB disease is more likely to reflect low frequency of persistent viable (“reactivate-able”) infection than low frequency of breakout of *Mtb* replication from long-term immunological control. The empirical evidence that we present in support of this contention is consistent with recent re-evaluations of epidemiological data which suggest that (1) duration of *Mtb* infection viability is likely to be much shorter than previously believed [20] and that (2) reactivation rates in IGRA or TST positive individuals unprotected by PT undergoing immunosuppressive therapy are much lower than would be expected if such testing represented infection truly capable of reactivation [21]. Emerging mathematical modelling outputs add weight to this paradigm shift, suggesting that a significant proportion of *Mtb*-infected individuals achieve self-clearance, leaving a much smaller population with persisting viable *Mtb* infection than previously assumed [22]. Finally, a precedent for lasting anti-mycobacterial immunological reactivity in the absence of bacterial viability already exists in the form of erythema nodosum leprosum, type II reactions to persistent *M. leprae* antigens which are known to occur years after mycobacterial cure.

These blood transcriptional responses to PT suggest that around one third of our IGRA+ study participants had true (viable) LTBI. This proportion is predicted to be lower with increasing remoteness in time since exposure [20]. The implications for national and global estimates of LTBI prevalence that rely upon IGRA/ TST data are clear and suggest a large overestimation of the size of the global reservoir of potentially reactivatable latent infection; we contend that such data should in future be presented as prevalence of tuberculin sensitivity and that the term LTBI should be used more judiciously. Since all incident reactivation arises from the true LTBI pool, the incidence rate in this subgroup of all IGRA positives will be considerably higher than, for example, the 0.6 per 100 person-years seen in the placebo arm of a recent vaccine trial [23]. The development of tools and strategies to readily identify this true LTBI subgroup would facilitate more efficient targeting of interventions to interrupt reactivation and would accelerate evaluation of novel interventions because the sample size required for future vaccine trials and trials of preventive therapy would be considerably reduced. Evaluations of risk factors associated with infection, premised on the use of IGRA/TST to define infection, have likely been using a very imperfect endpoint with the associated high likelihood of misclassification error.

The temporal dynamics of the transcriptomic changes are such that evidence of a response can be detected as early as 2 weeks into PT. This raises the possibility of a ‘treat and test’ approach to PT wherein the absence of a specific change in a biomarker (or biomarker profile) at an early time point, say 2 weeks into treatment, could be interpreted as an indication that further treatment will have no effect and can then be discontinued. Recent TB host gene expression studies have shown that biomarker signatures can be shrunk to small sets with the potential to be implemented as diagnostic or prognostic tests in the field [24–26].

This study had a relatively modest number of participants. Our results based on the transcriptional profiles after PT therapy should be studied in larger prospective cohorts with well-defined clinical outcomes and long term follow up. Sequential transcriptomic and cell count differential testing on a larger study population (including children), with a variety of exposure histories and diverse PT regimens (including those under investigation for multidrug-resistant LTBI) will help to elucidate the array of responses encountered. The hunt for predictors of future disease amongst TB-exposed individuals has previously been directed towards identification of biomarkers indicating increased risk, an approach that risks dismissal of future changes in the host environment which it might not be possible to anticipate (e.g. transplant immunosuppression). By removing from the pool of *Mtb*-sensitized participants (IGRA+ or TST+) a significant proportion for whom reactivation is biologically impossible (because no viable *Mtb* infection remains), the scale of the prevention challenge is drastically reduced and a more efficient targeted and nuanced approach can be considered.

Validation of this transcriptomic signature in ongoing trials of PT in which defined secondary cases are identified is now a priority. Important implications of a test that can distinguish IGRA+ or TST+ *Mtb* sensitized individuals at zero risk of progression/reactivation include drastic reevaluation of the global burden of LTBI, stratification of preventive therapy and post-exposure vaccine efficacy, higher resolution targeting of LTBI preventive therapy, potential use as a biomarker for efficacy evaluation of novel PT regimens for drug-susceptible and drug-resistant-TB, and PT test of cure.

Individuals with immunological memory of a prior encounter with *Mtb* (commonly referred to as LTBI) who are treated with PT demonstrate two different phenotypes of transcriptomic response. We propose that the clear responders are those who had truly viable latent *Mtb* infection, and that the minimal responders, in common with the IGRA-negative, previously unexposed healthy controls, had no viable *Mtb* organisms and were therefore not truly latently TB infected.

## Materials and Methods

### Participants

Study participants were recruited from National Health Service (NHS) tuberculosis (TB) outpatient clinics in London (Whittington Health NHS Trust, Royal Free London NHS Foundation Trust, Barts Health NHS Trust, Homerton University Hospital NHS Foundation Trust). Healthy controls were recruited from the London School of Hygiene and Tropical Medicine.

Participants were recruited who were aged 18 years and above, had positive Interferon Gamma Release Assay (IGRA) (performed by the local hospital laboratories, using the QuantiFERON-TB Gold In-tube assay [Qiagen, Manchester, UK]), with known exposure to an index person with isoniazid and rifampicin susceptible pulmonary TB (unconfirmed for three individuals) and who planned to initiate a 12-week course of combined rifampicin/ isoniazid (RH) as preventive therapy (once daily rifampicin 600 mg/ isoniazid 300 mg as Rifinah) plus once daily pyridoxine 10 mg. Adult volunteers aged 18 years and above were recruited as healthy control participants.

Once consented, demographic information, TB exposure history, and medical history were recorded on a data capture sheet and testing for human immunodeficiency virus (HIV) was performed. Healthy volunteers additionally underwent IGRA testing (performed using the QuantiFERON-TB Gold In-tube assay according to the manufacturer’s recommendations) and were excluded if they were found to be IGRA+. Individuals were excluded if they had a prior history of TB infection, of having taken anti-TB treatment or exposure to drug-resistant TB. Participants who were pregnant, breastfeeding or trying to conceive, those with immunosuppressive disorders including HIV and those who had taken immunosuppressant medication in the preceding six months were also excluded. Healthy control participants reporting prior exposure to TB were also excluded.

Healthy controls were given a two-week course of RH (once daily rifampicin 600 mg/ isoniazid 300 mg as Rifinah) plus once daily pyridoxine 10 mg.

Blood samples were collected from all participants at baseline (V1) and 2 weeks after initiating RH (V2), with an additional sample point in IGRA+ participants within 6 weeks of completion of the 12-week course of treatment (V3). At all sampling timepoints, all participants were asked about their adherence to treatment, and 2.5 ml whole blood was collected in a PAXgene blood RNA tube (PreAnalytiX GmbH, Hombrechtikon, Switzerland) for RNA expression analysis and a Lithium heparin tube (Becton Dickinson, Berkshire, UK) for subsequent stimulation assays. The PAXgene tubes were frozen within 4 hours of collection.

The study procedures and protocol were approved by City & East NHS Research Ethics Committee, London (reference 16/LO/1206) and the London School of Hygiene and Tropical Medicine Research Ethics Committee (reference 11603). Written informed consent was given by all participants before inclusion in the study.

### Stimulation of whole blood

Stimulation was performed using QuantiFERON-TB Gold Plus In-tube Assay (QFT-TB Plus) (Qiagen). Within four hours of collection, 1 ml of blood was transferred from the lithium heparin tube to each of the four QFT-TB Plus tubes – TB1 antigen, TB2 antigen (both containing peptides from ESAT-6 and CFP-10 antigens), mitogen positive control and (unstimulated) negative control – the tubes were gently shaken to dissolve the lyophilized peptides in the blood. The QFT-TB Plus tubes were immediately incubated upright at 37°C for 22 −24 hours. After incubation, the blood was transferred into a 1.5 ml microcentrifuge tube and centrifuged for 15 minutes at 3000 RCF(g). Supernatants were removed and the remaining cell pellet (500 μl) was transferred into a 15 ml tube containing 2.5 ml RNAprotect^®^ Cell Reagent (Qiagen). The cells were resuspended by vortexing, and incubated for 2 hours for complete cell lysis before freezing at −80°C.

### Peripheral blood RNA expression by microarray

Total RNA was extracted from the PAXgene tubes using the PAXgene Blood miRNA Kit (Qiagen), and from the QFT-TB Plus stimulated samples, which had been lysed in RNAprotect, using the RNEasy mini kit (Qiagen), according to the manufacturer’s instructions, incorporating on-column DNAse digestion. Globin depletion was performed using the GLOBINclear Kit (ThermoFisher), quantified by Nanodrop and the quality was assessed using an Agilent Bioanalyzer (Agilent, Cheshire, UK. The two-color low input Quick Amp Labelling kit (Agilent) was used to Cy3- or Cy5-fluorescently label cRNA samples, which were then hybridized to SurePrint G3 Human Gene Expression 60K GeneChip microarrays (Agilent) according to the manufacturer’s instructions. Hybridization intensity was quantified via a SureScan Microarray Scanner (Agilent). Microarray data are deposited at Gene Expression Omnibus, Series GSE153342.

Individual channel intensities from the GeneChip data were extracted independently and analysed as separate observations [27].

### Statistical analyses

Clinical data were analysed using ‘R’ Language and Environment for Statistical Computing 3.5.2. Fishers, Chi-squared and Kruskall Wallis tests of significance were used for categorical data. Mann-Whitney U tests of significance were used for continuous data.

Expression data were analysed using ‘R’ Language and Environment for Statistical Computing 3.5.2. Pre-processing, log-2 transformation and normalisation were performed using the Agilp package [28]. Microarrays were run using two batches of microarray slides and Principal Component Analysis identified an associated batch effect. Batch correction was performed using the COmBat function in the Surrogate Variable Analysis (sva) package in R [29, 30]. To minimise the potential influence of batch correction on subsequent clustering analyses, no reference batch was used and independent COmBat-corrections were performed for each dataset of interest (individual PAXgene, TB1 and TB2 tube datasets and a combined TB1/ TB2/ negative tube dataset). Post-Combat correction PCA plots were undertaken to confirm the removal of the batch effect and identify outliers (Figure 1– Figure supplements 1 and 2).

Differential gene expression analysis was performed using the limma package in R [4] which uses linear models. Where paired samples were available and analysis was relevant, paired t-tests were performed, with this being stated in the results. Adjustment for false discovery rate was performed using Benjamini-Hochberg (BH) correction with a significance level of adjusted p-value <0.05.

Prior to longitudinal analyses, the gene expression set was filtered to remove noise. Lowly expressed transcripts for which expression values did not exceed a value of 6 for any of the samples, were removed. Transcripts with extreme outlying values were removed, which were defined as values < (Quartile1 – [3* Inter-Quartile Range]) or > (Quartile3 + [3 * Inter-Quartile Range]). Transcripts with the greatest temporal and interpersonal variability were then selected based on their variance, with those transcripts with variance >0.1 taken forwards to the longitudinal analysis. X-chromosome transcripts which were significantly differentially expressed with gender at V1, V2 and/ or V3 were identified using linear models in limma (BH corrected p value < 0.05) and were excluded, as were Y-chromosome transcripts.

Unsupervised longitudinal clustering analyses were performed using the BClustLong package in ‘R’ [31], which uses a Dirichlet process mixed model for clustering longitudinal gene expression data. A linear mixed-effects framework is used to model the trajectory of genes over time and bases clustering on the regression coefficients obtained from all genes. 500 iterations were run (thinning by 2, so 1000 iterations in total).

Longitudinal differential gene expression analyses were performed using the MaSigPro package in R [7]. MaSigPro follows a two steps regression strategy to find genes with significant temporal expression changes and significant differences between groups. Coefficients obtained in the second regression model are used to then cluster together significant genes with similar expression patterns. Adjustment for false discovery rate was performed using BH correction with a significance level of adjusted p-value <0.05. Given the three timepoints from the IGRA+ individuals and the two timepoints from the healthy control groups, we employed both quadratic and linear approaches to account for all the potential curve shapes in the gene expression data.

Estimations of relative cellular abundances were calculated from the normalised full gene expression matrix (58,201 gene probes) using the CibersortX [16], which uses gene expression data to deconvolve mixed cell populations. We used the LM22 [32] leukocyte gene signature matrix as reference, that comprises 22 different immune cell types, and ran 1,000 permutations. Total monocyte fraction was calculated as the sum of the fractions of monocytes, macrophages and dendritic cells. Total lymphocyte fraction was calculated as the sum of B cells, Plasma cells, CD8+ T cells, CD4+ T cells, Helper follicular T cells, Regulatory T cells, Gamma delta T cells, and NK cells. A polynomial model (degrees of freedom = 2) was fitted in R to estimate relationships between the monocyte: lymphocyte ratio and time, in IGRA+ subgroups A and B.

## Acknowledgements

The authors wish to thank the patients and volunteers who participated in the study. We also thank the clinical staff at Barts Health NHS Trust, Homerton University Hospital Foundation Trust and TB Service North Central London, in particular Heinke Kunst (Barts Health NHS Trust), Graham Bothamley (Homerton University Hospital Foundation Trust) and Marc Lipman (TB Service North Central London). The authors also wish to thank the research nurses who assisted with this study, including Victoria Dean and Michelle Berin (University College London) and Nirmala Ghimire (Barts Health) and Ortensia Vito (Imperial College London) for help with data analysis.

This work was funded by a British Infection Association Small Project Research Grant (2016) and a Rosetrees Trust Seed Corn Award (# JS15 / M660). C.B. was funded by an Academic Clinical Fellowship from the National Institute for Health Research (NIHR) (ACF-2012-18-008) and currently receives support from an Imperial 4i Wellcome Trust/ NIHR Imperial BRC Clinical PhD Fellowship. M.K. receives support from the NIHR Imperial College BRC and the Wellcome Trust (Sir Henry Wellcome Fellowship grant no. 206508/Z/17/Z). JC receives support from the Medical Research Council Newton Fund (*#*MR/P017568/1).

## Competing Interests

No competing interests are declared by the authors.

## Supplementary Data

### Supplementary Figures

**Figure 1- figure supplement 1.**
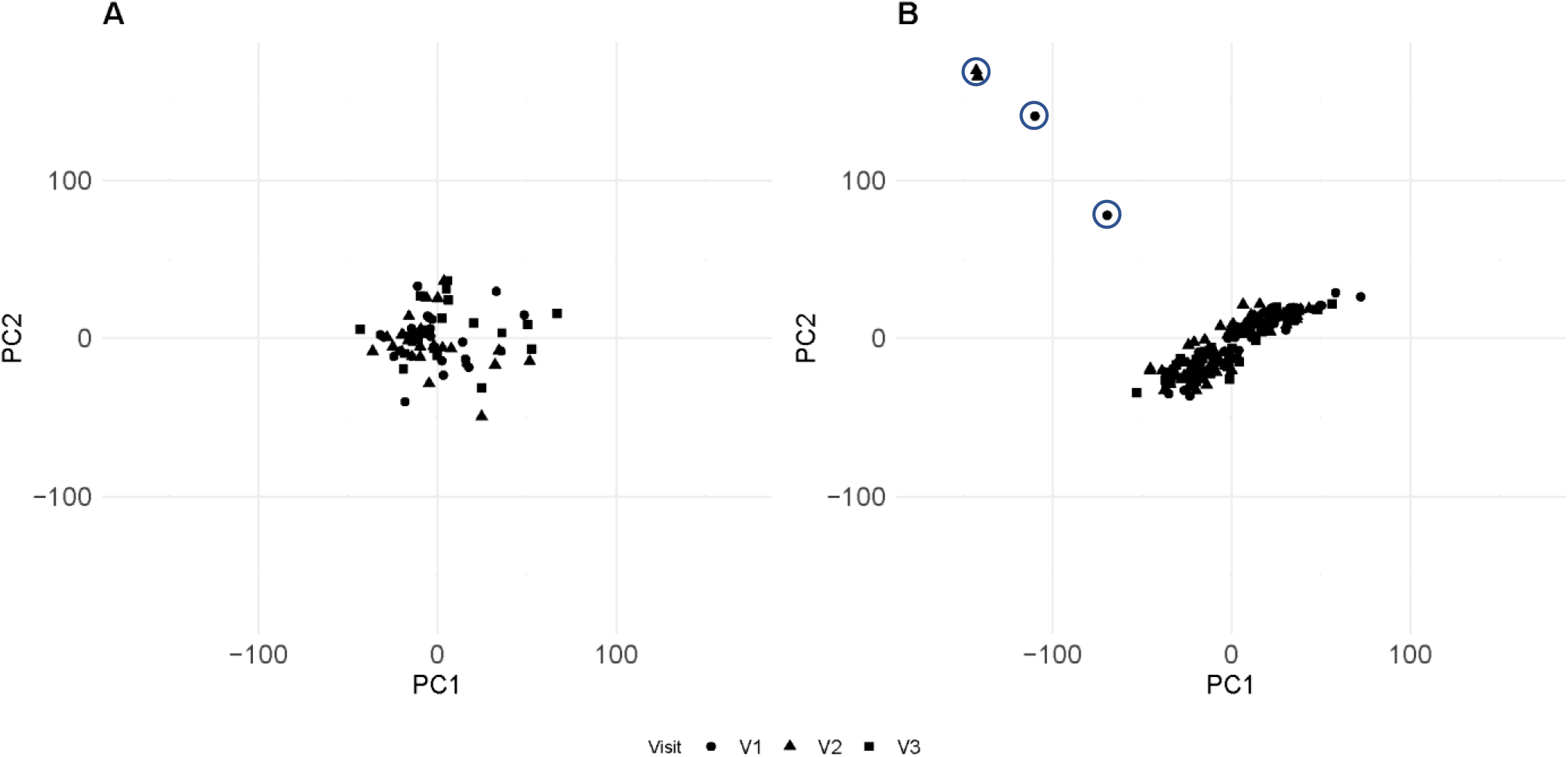
Principle component analyses (PCA) of the gene expression sets were performed. Plots showing dimensions 1 and 2 of the PCA of the PAXgene samples (A) and the stimulated samples (B) before ComBat correction. In the stimulated samples, healthy control (HC52) was an outlier in dimensions 1 and 2 (circled) and this persisted after batch correction (not shown), so HC52 was excluded from the subsequent analyses.

**Figure 1- figure supplement 2.**
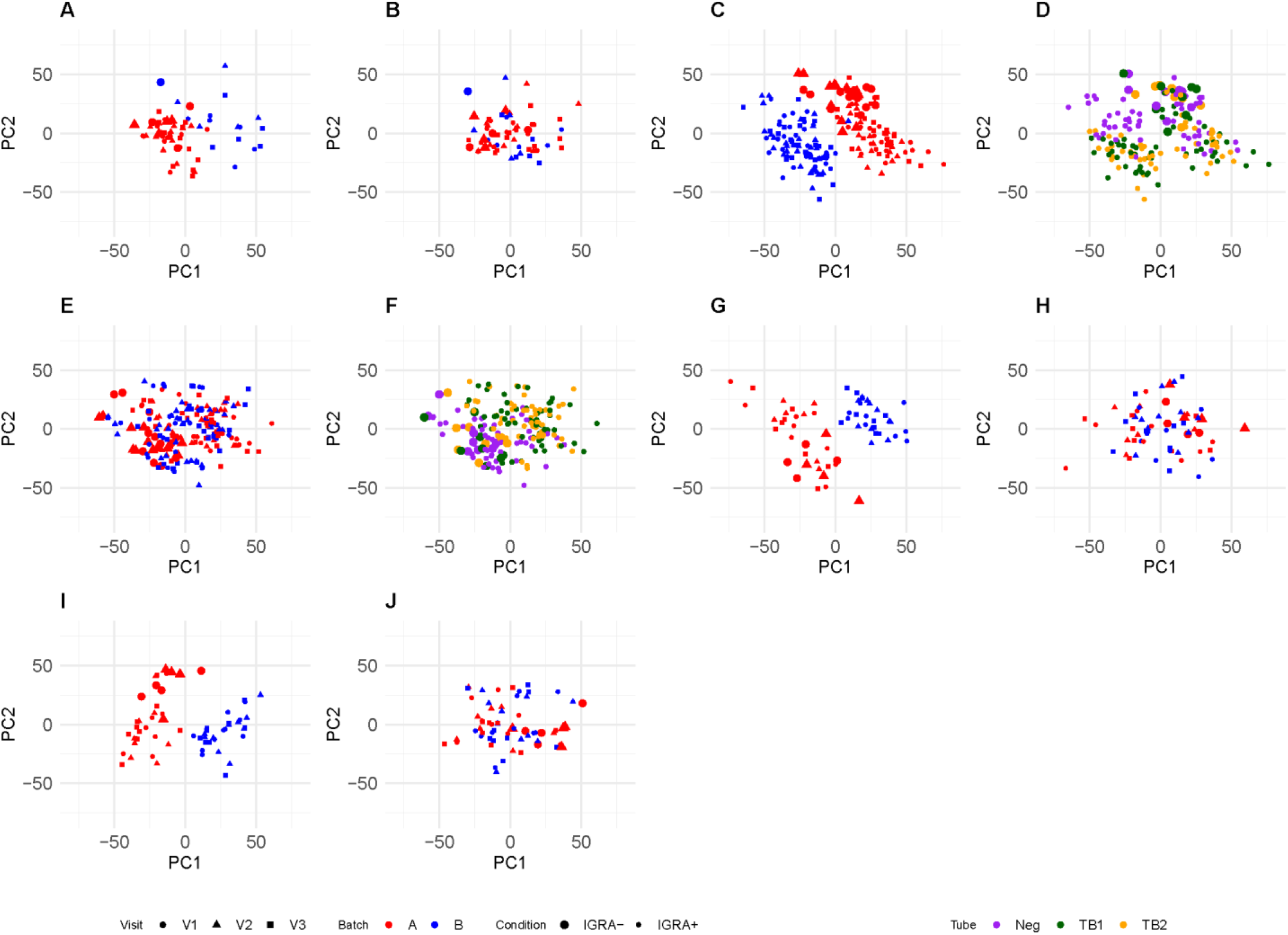
Gene expression data from 18 IGRA+ and 4 IGRA-participants were included in the final analyses. Principle component analyses (PCA) of the gene expression sets were performed before and after batch correction with ComBat. Plots showing dimensions 1 and 2 of the PCA of the PAXgene tube samples before (A) and after ComBat (B); all stimulated samples (TB1, TB2 and Negative) before (C, D) and after ComBat (E, F) with C and E showing batch differentiation and D and F showing tube differentiation; TB1 samples before (G) and after Combat (H); TB2 samples before (I) and after Combat (J). Batch, visit, IGRA status and QuantiFERON TB Gold plus tube are provided.

**Figure 2- figure supplement 1.**
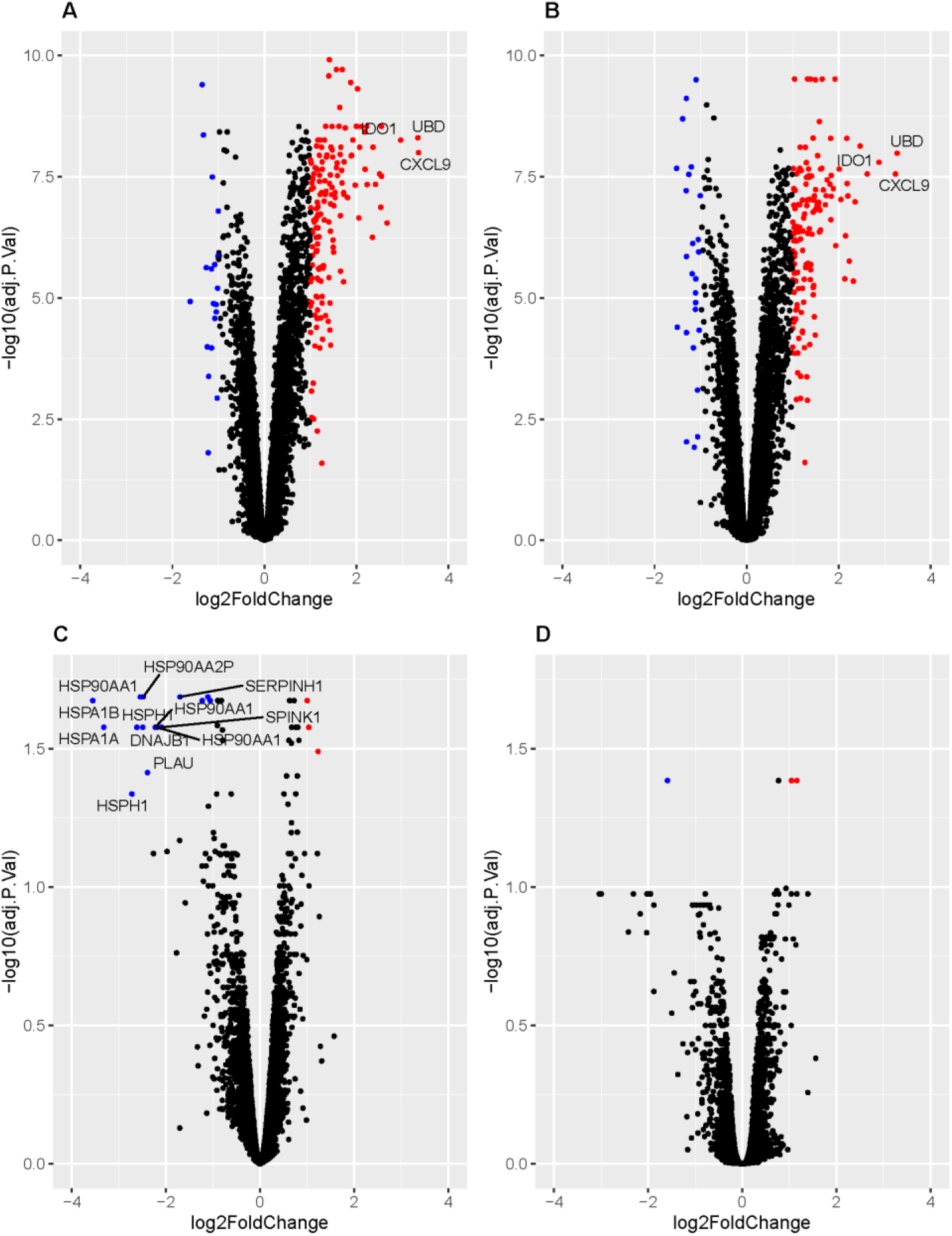
Volcano plots showing genes significantly differentially expressed between stimulated (QuantiFERON Gold Plus TB1 and TB2 tubes) and unstimulated (QuantiFERON Gold Plus negative tubes) blood samples. Genes overexpressed in stimulated blood with log2Foldchange (LFC) >1 and BH adjusted p value <0.05 are shown in red. Genes underexpressed in stimulated blood with LFC <-1 and BH adjusted p value <0.05 are shown in blue. Genes with LFC >2.7 and < −1.7 are annotated with their gene symbols. Plots are shown for IGRA+ subjects, comparing TB1 *vs*. negative tube samples (A), and TB2 *vs*. negative tube samples (B) at visit 1. Also shown are plots for IGRA-subjects, comparing TB1 *vs*. negative tube samples (C), and TB2 *vs*. negative tube samples (D) at visit 1.

**Figure 4 - figure supplement 1.**
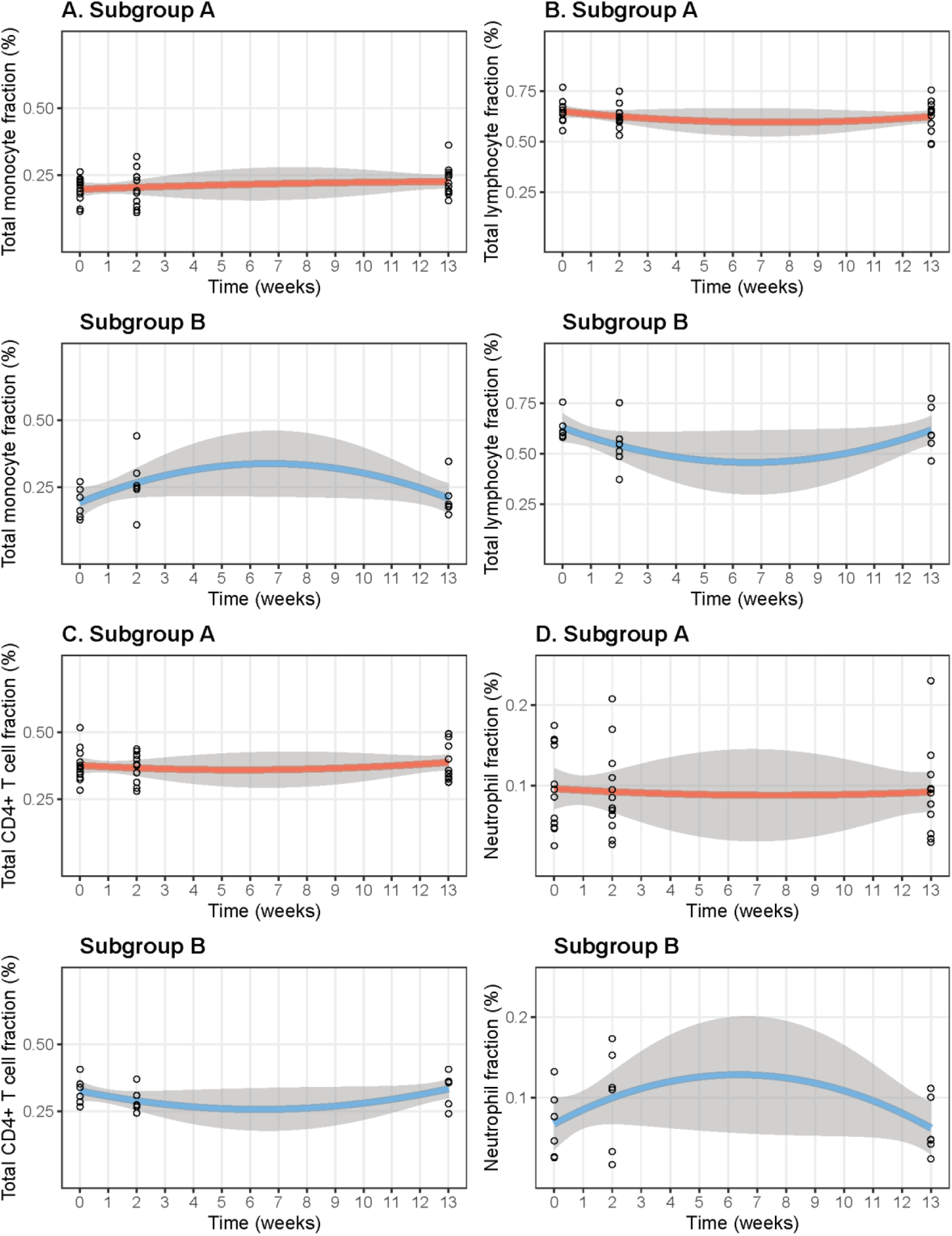
Cibersortx was used to estimate the abundance of different cell types in the TB2-stimulated whole blood samples at each visit. Scatterplots showing the change cellular fractions over the time-course of the study period in IGRA+ subgroups A and B for Total monocyte fraction (A), Total lymphocyte fraction (B), Total CD4+ T cell fraction (C), Neutrophil fraction (D). Visit 1 is 0 weeks, Visit 2 is 2 weeks and Visit 3 is 13 weeks, with 90% confidence intervals shown.

### Supplementary Files

Supplementary File 1: Significantly differentially expressed (SDE) transcripts IGRA+ vs IGRA-in TB1 tubes at Visit (V) 1 and V2 and in TB2 tubes at V1 and V2.

Supplementary File 2: SDE transcripts TB1 *vs* negative tube, TB2 *vs* negative tube at V1, in IGRA+.

Supplementary File 3: SDE transcripts TB1 *vs* negative tube, TB2 *vs* negative tube at V1, in IGRA-.

Supplementary File 4: 474 most variable transcripts (TB2-stimulated samples).

Supplementary File 5: MaSigPro results: transcripts SDE though time, Cluster 1 *vs* Cluster 2.

Supplementary File 6: Results of biological pathways analysis using Reactome pathway knowledgebase.

## Notes

### Competing Interest Statement

The authors have declared no competing interest.

## References

1. World Health Organisation. Latent tuberculosis infection: updated and consolidated guidelines for programmatic management. 2018: Geneva.

2. Esmail, H., C.E. Barry, 3rd, D.B. Young, and R.J. Wilkinson, The ongoing challenge of latent tuberculosis. Philos Trans R Soc Lond B Biol Sci, 2014. 369(1645): p. 20130437.

3. Whalen, C.C., J.L. Johnson, A. Okwera, D.L. Hom, R. Huebner, P. Mugyenyi, R.D. Mugerwa, and J.J. Ellner, A trial of three regimens to prevent tuberculosis in Ugandan adults infected with the human immunodeficiency virus. Uganda-Case Western Reserve University Research Collaboration. N Engl J Med, 1997. 337(12): p. 801–8.

4. Ritchie, M.E., B. Phipson, D. Wu, Y. Hu, C.W. Law, W. Shi, and G.K. Smyth, limma powers differential expression analyses for RNA-sequencing and microarray studies. Nucleic Acids Res, 2015. 43(7): p. e47.

5. Petruccioli, E., T. Chiacchio, I. Pepponi, V. Vanini, R. Urso, G. Cuzzi, L. Barcellini, D.M. Cirillo, F. Palmieri, G. Ippolito, and D. Goletti, First characterization of the CD4 and CD8 T-cell responses to QuantiFERON-TB Plus. J Infect, 2016. 73(6): p. 588–597.

6. Sun, J., J.D. Herazo-Maya, N. Kaminski, H. Zhao, and J.L. Warren, A Dirichlet process mixture model for clustering longitudinal gene expression data. Stat Med, 2017. 36(22): p. 3495–3506.

7. Conesa, A. and M.J. Nueda. maSigPro: Significant Gene Expression Profile Differences in Time Course Gene Expression Data. R package version 1.54.0. 2018; Available from: http://bioinfo.cipf.es/.

8. Anderson, S.T., M. Kaforou, A.J. Brent, V.J. Wright, C.M. Banwell, G. Chagaluka, A.C. Crampin, H.M. Dockrell, N. French, M.S. Hamilton, M.L. Hibberd, F. Kern, P.R. Langford, L. Ling, R. Mlotha, T.H. Ottenhoff, S. Pienaar, V. Pillay, J.A. Scott, H. Twahir, R.J. Wilkinson, L.J. Coin, R.S. Heyderman, M. Levin, B. Eley, I. Consortium, and K.T.S. Group, Diagnosis of childhood tuberculosis and host RNA expression in Africa. N Engl J Med, 2014. 370(18): p. 1712–23.

9. Berry, M.P., C.M. Graham, F.W. McNab, Z. Xu, S.A. Bloch, T. Oni, K.A. Wilkinson, R. Banchereau, J. Skinner, R.J. Wilkinson, C. Quinn, D. Blankenship, R. Dhawan, J.J. Cush, A. Mejias, O. Ramilo, O.M. Kon, V. Pascual, J. Banchereau, D. Chaussabel, and A. O’Garra, An interferon-inducible neutrophil-driven blood transcriptional signature in human tuberculosis. Nature, 2010. 466(7309): p. 973–7.

10. Blankley, S., C.M. Graham, J. Turner, M.P. Berry, C.I. Bloom, Z. Xu, V. Pascual, J. Banchereau, D. Chaussabel, R. Breen, G. Santis, D.M. Blankenship, M. Lipman, and A. O’Garra, The Transcriptional Signature of Active Tuberculosis Reflects Symptom Status in Extra-Pulmonary and Pulmonary Tuberculosis. PLoS One, 2016. 11(10): p. e0162220.

11. Bloom, C.I., C.M. Graham, M.P. Berry, F. Rozakeas, P.S. Redford, Y. Wang, Z. Xu, K.A. Wilkinson, R.J. Wilkinson, Y. Kendrick, G. Devouassoux, T. Ferry, M. Miyara, D. Bouvry, V. Dominique, G. Gorochov, D. Blankenship, M. Saadatian, P. Vanhems, H. Beynon, R. Vancheeswaran, M. Wickremasinghe, D. Chaussabel, J. Banchereau, V. Pascual, L.P. Ho, M. Lipman, and A. O’Garra, Transcriptional blood signatures distinguish pulmonary tuberculosis, pulmonary sarcoidosis, pneumonias and lung cancers. PLoS One, 2013. 8(8): p. e70630.

12. Kaforou, M., V.J. Wright, T. Oni, N. French, S.T. Anderson, N. Bangani, C.M. Banwell, A.J. Brent, A.C. Crampin, H.M. Dockrell, B. Eley, R.S. Heyderman, M.L. Hibberd, F. Kern, P.R. Langford, L. Ling, M. Mendelson, T.H. Ottenhoff, F. Zgambo, R.J. Wilkinson, L.J. Coin, and M. Levin, Detection of tuberculosis in HIV-infected and -uninfected African adults using whole blood RNA expression signatures: a case-control study. PLoS Med, 2013. 10(10): p. e1001538.

13. Maertzdorf, J., J. Weiner, 3rd, H.J. Mollenkopf, T.B. Network, T. Bauer, A. Prasse, J. Muller-Quernheim, and S.H. Kaufmann, Common patterns and disease-related signatures in tuberculosis and sarcoidosis. Proc Natl Acad Sci U S A, 2012. 109(20): p. 7853–8.

14. Ottenhoff, T.H., R.H. Dass, N. Yang, M.M. Zhang, H.E. Wong, E. Sahiratmadja, C.C. Khor, B. Alisjahbana, R. van Crevel, S. Marzuki, M. Seielstad, E. van de Vosse, and M.L. Hibberd, Genome-wide expression profiling identifies type 1 interferon response pathways in active tuberculosis. PLoS One, 2012. 7(9): p. e45839.

15. Jassal, B., L. Matthews, G. Viteri, C. Gong, P. Lorente, A. Fabregat, K. Sidiropoulos, J. Cook, M. Gillespie, R. Haw, F. Loney, B. May, M. Milacic, K. Rothfels, C. Sevilla, V. Shamovsky, S. Shorser, T. Varusai, J. Weiser, G. Wu, L. Stein, H. Hermjakob, and P. D’Eustachio, The reactome pathway knowledgebase. Nucleic Acids Res, 2020. 48(D1): p. D498–D503.

16. Newman, A.M., C.B. Steen, C.L. Liu, A.J. Gentles, A.A. Chaudhuri, F. Scherer, M.S. Khodadoust, M.S. Esfahani, B.A. Luca, D. Steiner, M. Diehn, and A.A. Alizadeh, Determining cell type abundance and expression from bulk tissues with digital cytometry. Nat Biotechnol, 2019. 37(7): p. 773–782.

17. Bloom, C.I., C.M. Graham, M.P. Berry, K.A. Wilkinson, T. Oni, F. Rozakeas, Z. Xu, J. Rossello-Urgell, D. Chaussabel, J. Banchereau, V. Pascual, M. Lipman, R.J. Wilkinson, and A. O’Garra, Detectable changes in the blood transcriptome are present after two weeks of antituberculosis therapy. PLoS One, 2012. 7(10): p. e46191.

18. Cliff, J.M., J.S. Lee, N. Constantinou, J.E. Cho, T.G. Clark, K. Ronacher, E.C. King, P.T. Lukey, K. Duncan, P.D. Van Helden, G. Walzl, and H.M. Dockrell, Distinct phases of blood gene expression pattern through tuberculosis treatment reflect modulation of the humoral immune response. J Infect Dis, 2013. 207(1): p. 18–29.

19. Naranbhai, V., H.A. Fletcher, R. Tanner, M.K. O’Shea, H. McShane, B.P. Fairfax, J.C. Knight, and A.V. Hill, Distinct Transcriptional and Anti-Mycobacterial Profiles of Peripheral Blood Monocytes Dependent on the Ratio of Monocytes: Lymphocytes. EBioMedicine, 2015. 2(11): p. 1619–26.

20. Behr, M.A., P.H. Edelstein, and L. Ramakrishnan, Revisiting the timetable of tuberculosis. BMJ, 2018. 362: p. k2738.

21. Behr, M.A., P.H. Edelstein, and L. Ramakrishnan, Is Mycobacterium tuberculosis infection life long? BMJ, 2019. 367: p. l5770.

22. Emery, J.C., A.S. Richards, K.D. Dale, C.F. McQuaid, R.G. White, D.J.T., and R.M.G.J. Houben. Self-clearance of Mycobacterium tuberculosis infection: implications for lifetime risk and population at-risk of tuberculosis disease [Poster presentation]. 50th Union World Conference on Lung Health, 2019 Oct 30-Nov 2. Hyderabad, India.

23. Tait, D.R., M. Hatherill, O. Van Der Meeren, A.M. Ginsberg, E. Van Brakel, B. Salaun, T.J. Scriba, E.J. Akite, H.M. Ayles, A. Bollaerts, M.A. Demoitie, A. Diacon, T.G. Evans, P. Gillard, E. Hellstrom, J.C. Innes, M. Lempicki, M. Malahleha, N. Martinson, D. Mesia Vela, M. Muyoyeta, V. Nduba, T.G. Pascal, M. Tameris, F. Thienemann, R.J. Wilkinson, and F. Roman, Final Analysis of a Trial of M72/AS01E Vaccine to Prevent Tuberculosis. N Engl J Med, 2019. 381(25): p. 2429–2439.

24. Roe, J.K., N. Thomas, E. Gil, K. Best, E. Tsaliki, S. Morris-Jones, S. Stafford, N. Simpson, K.D. Witt, B. Chain, R.F. Miller, A. Martineau, and M. Noursadeghi, Blood transcriptomic diagnosis of pulmonary and extrapulmonary tuberculosis. JCI Insight, 2016. 1(16): p. e87238.

25. Suliman, S., E. Thompson, J. Sutherland, J. Weiner Rd, M.O.C. Ota, S. Shankar, A. Penn-Nicholson, B. Thiel, M. Erasmus, J. Maertzdorf, F.J. Duffy, P.C. Hill, E.J. Hughes, K. Stanley, K. Downing, M.L. Fisher, J. Valvo, S.K. Parida, G. van der Spuy, G. Tromp, I.M.O. Adetifa, S. Donkor, R. Howe, H. Mayanja-Kizza, W.H. Boom, H. Dockrell, T.H.M. Ottenhoff, M. Hatherill, A. Aderem, W.A. Hanekom, T.J. Scriba, S.H. Kaufmann, D.E. Zak, G. Walzl, G.C. and the, and A.C.S.c.s. groups, Four-gene Pan-African Blood Signature Predicts Progression to Tuberculosis. Am J Respir Crit Care Med, 2018.

26. Sweeney, T.E., L. Braviak, C.M. Tato, and P. Khatri, Genome-wide expression for diagnosis of pulmonary tuberculosis: a multicohort analysis. Lancet Respir Med, 2016. 4(3): p. 213–24.

27. Chain, B., H. Bowen, J. Hammond, W. Posch, J. Rasaiyaah, J. Tsang, and M. Noursadeghi, Error, reproducibility and sensitivity: a pipeline for data processing of Agilent oligonucleotide expression arrays. BMC Bioinformatics, 2010. 11: p. 344.

28. Chain, B. agilp: Agilent expression array processing package. R package version 3.14.0. 2018; Available from: http://bioconductor.org/packages/release/bioc/html/agilp.html.

29. Johnson, W.E., C. Li, and A. Rabinovic, Adjusting batch effects in microarray expression data using empirical Bayes methods. Biostatistics, 2007. 8(1): p. 118–27.

30. Leek JT, Johnson WE, Parker HS, Fertig EJ, Jaffe AE, Storey JD, Zhang Y, and T. Lc. sva: Surrogate Variable Analysis. R package version 3.34.0. 2019; Available from: https://bioconductor.org/packages/release/bioc/html/sva.html.

31. Sun, J., J.D. Herazo-Maya, N. Kaminski, H. Zhao, and J.L. Warren. BClustLonG: A Dirichlet Process Mixture Model for Clustering Longitudinal Gene Expression Data. R package version 0.1.2. 2017; Available from: https://CRAN.R-project.org/package=BClustLonG.

32. Newman, A.M., C.L. Liu, M.R. Green, A.J. Gentles, W. Feng, Y. Xu, C.D. Hoang, M. Diehn, and A.A. Alizadeh, Robust enumeration of cell subsets from tissue expression profiles. Nat Methods, 2015. 12(5): p. 453–7.

